# Polypyrrole increases branching and neurite extension by Neuro2A cells on PBAT ultrathin fibers

**DOI:** 10.1101/241307

**Authors:** Alessandro E.C. Granato, André C. Ribeiro, Fernanda R. Marciano, Bruno V. M. Rodrigues, Anderson O. Lobo, Marimelia Porcionatto

## Abstract

**Graphical Abstract:** 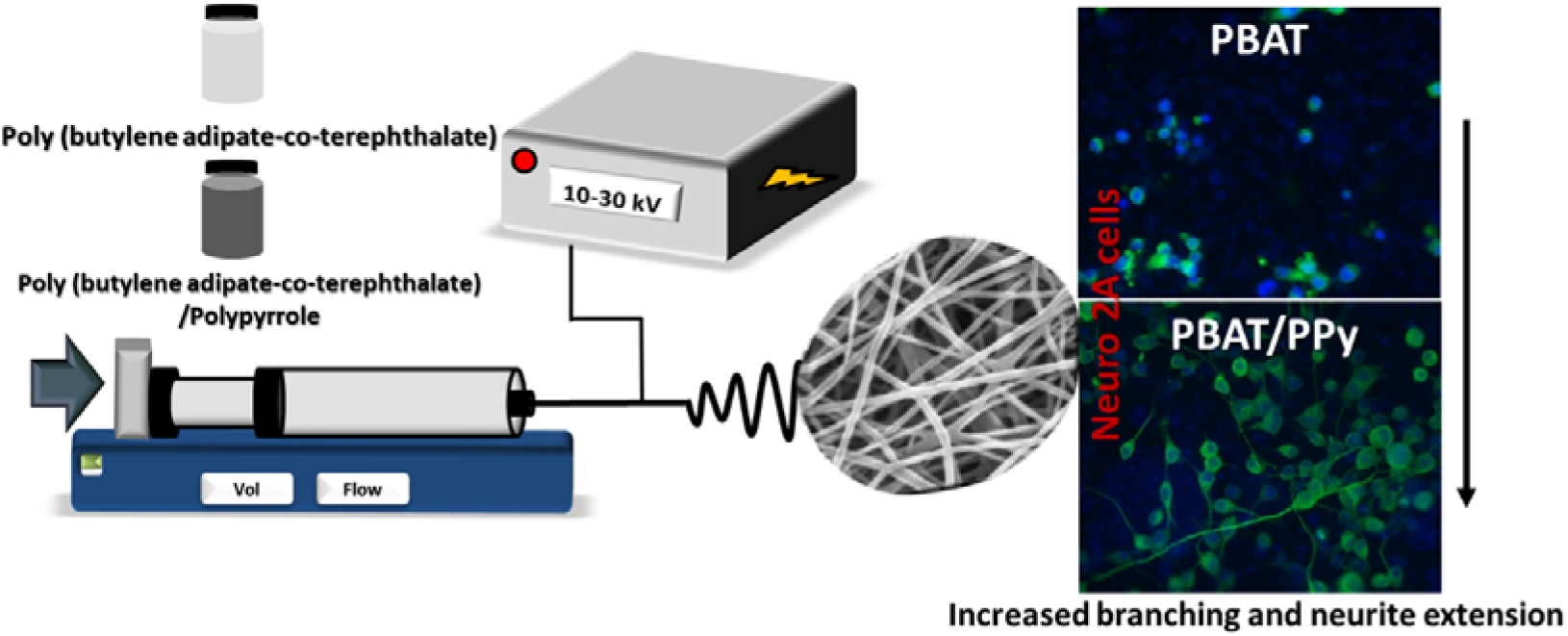

Electrospinning shows a feasible way to generate hybrid scaffolds from the combination of different materials. This work presented a successful route to prepare ultrathin fibers from hybrid solutions containing a commercial polyester, poly (butylene adipate-co-terephthalate) (PBAT) and a conductive polymer, polypyrrole (PPy). The final material (PBAT/PPy) showed an enhanced potential for neuronal differentiation when compared to neat PBAT. The PPy loading improved branching and neurite extension of Neuro2a cells, which opens a wide range of perspectives where these materials may be applied in regenerative medicine.

**ABSTRACT:** We present a methodology for production and application of electrospun hybrid materials containing commercial polyester (poly (butylene adipate-co-terephthalate; PBAT), and a conductive polymer (polypirrole; PPy) as scaffold for neuronal growth and differentiation. The physical-chemical properties of the scaffolds and optimization of the electrospinning parameters are presented. The electrospun scaffolds are biocompatible and allow proper adhesion and spread of mesenchymal stem cells (MSCs). Fibers produced with PBAT with or without PPy were used as scaffold for Neuro2a mouse neuroblastoma cells adhesion and differentiation. Neuro2a adhered to PBAT and PBAT/PPy2% scaffolds without laminin coating. However, Neuro2a failed to differentiate in PBAT when stimulated by treatment with retinoic acid (RA), but differentiated in PBAT/PPy2% fibers. We hypothesize that PBAT hydrophobicity inhibited proper spreading and further differentiation, and inhibition was overcome by coating the PBAT fibers with laminin. We conclude that fibers produced with the combination of PBAT and PPy can support neuronal differentiation.

## BACKGROUND

The growing prevalence of neurodegenerative diseases, in addition to the occurrence of injuries to the central nervous system involving complex regenerative processes, has made the development of materials that may favor the growth and differentiation of neural cells one of the major challenges of regenerative medicine. Along with the greater understanding of cellular interactions, more attention has been given to the development of frameworks that are able to stimulate these regenerative processes. While cellular mechanisms can vary greatly in different tissue types, all engineered materials must ideally mimic the environment of the natural tissue to meet the mechanical, structural and functional properties of the natural tissue (1).

In the last two decades, electrospinning has emerged as a powerful tool in tissue engineering, due to its versatility in the production of micro, ultrathin (100 nm < diameter <1 000 nm), and nanofibers (diameter < 100 nm) (2, 3). Electrospinning uses a high electric field to produce ultrafine polymeric fibers with diameters ranging from 2 nm to several micrometers using natural and synthetic polymers (2, 4). This process offers unique capabilities for producing novel nanofibers and fabrics with controllable pore structure (5).

Because of this particular range of dimensions, electrospun fibers have been used as platforms to mimic the extracellular matrix (1). For instance, fibers produced by electrospinning may present notable lengths and uniform diameter both arising from the elongation of viscoelastic jets from a polymer solution or melt.

Conductive polymers (CP) have equivalent electrical properties compared to those of semiconductors, or even some metals, which provide them the ability to undergo electrical stimuli. These materials attracted much attention as potential candidates to be used as scaffolds since some of them present properties of biocompatibility, reversible conductivity by doping/dedoping, controllable chemical and electrochemical properties (6), easy processability and suitable wettability for cell adhesion (7).

Polypyrrole (PPy) is a CP with a widely recognized conductive property, which may be favorable in cell differentiation and growth. PPy allows for space-time control of electrical stimulation and generates minimal inflammatory response, which is beneficial for cell manipulation, regulatory approval, and clinical use (8). Studies using rodent stem cells, have demonstrated that PPy doped with bioactive molecules may influence cell survival and differentiation (9, 10).

Among the biodegradable polymers extensively studied in the last two decades, poly (butylene adipate-co-terephthalate) (PBAT), a commercially available and biodegradable co-polyester (11), has high potential for industrial and environmental applications. More recently, studies have been conducted considering the application of PBAT for biomedical purposes (12, 13, 14).

PBAT is biodegradable by microorganisms and has an adjustable balance between optimal biodegradation and desirable physical-chemical-properties (15). Thus, the use of PBAT, alone or in composite form, represents a great promise for the tissue engineering, since it presents desirable degradation rate, good thermal and mechanical properties and easy processability (16).

In this context, the electrospun PBAT/PPy ultrathin becomes of great interest, due to the combination of the biodegradability and biocompatibility of PBAT along with the conductive properties of PPy. Thus, through the capacity of electrical stimuli transmission of the material and its morphological similarity with the extracellular matrix, these biomaterials may present a high potential for application in the regeneration of nervous system.

To date, there is no study published using PBAT/PPy blends towards the use of neural cells for tissue engineering applications. Moreover, there is no study focusing on the effect of PPy in improving biocompatibility of hydrophobic co-polymers such as PBAT. We present for the first time electrospun PBAT/PPy fibers as scaffold for neuronal cell culture and differentiation. Our results show that PBAT fibers are biocompatible and when combined with PPy improve branching and neurite extension of Neuro2a cells. The results presented in this investigation suggest a promising potential of these materials for the future development of *in vivo* applications aiming at nervous tissue engineering e regeneration.

## MATERIAL AND METHODS

### Preparation of the polymer solutions

*N, N-*dimethylformamide (DMF, Sigma-Aldrich, ≥ 99%) and chloroform (Sigma-Aldrich, ≥ 99 %) were used as solvents. PPy was synthetized by chemical oxidation of pyrrole (BASF), using (NH_4_)_2_S_2_O_8_ as oxidizing agent (acid form) and purified by distillation, as described elsewhere (15).

In order to evaluate the best condition for the production of electrospun fibrous mats, different PBAT (commercial Ecoflex^®^ F Blend 107 C1200) concentrations were studied, viz. 8, 10, 12 and 14 wt %. Various solvent system ratios (chloroform:DMF) were also explored in order to obtain an optimum dispersion of PPy, viz. 50/50, 60/40, 75/25 and 85/15 (Table 1). PBAT was dissolved in chloroform (2 h, room temperature). Separately, PPy was homogeneously dispersed in DMF under sonication (VCX 500 – Sonics) to the concentration of 1–3 wt % during 60 min. Then, the different PBAT/PPy solutions were prepared and magnetically stirred for 20 h prior electrospinning process.

**Table 1:**
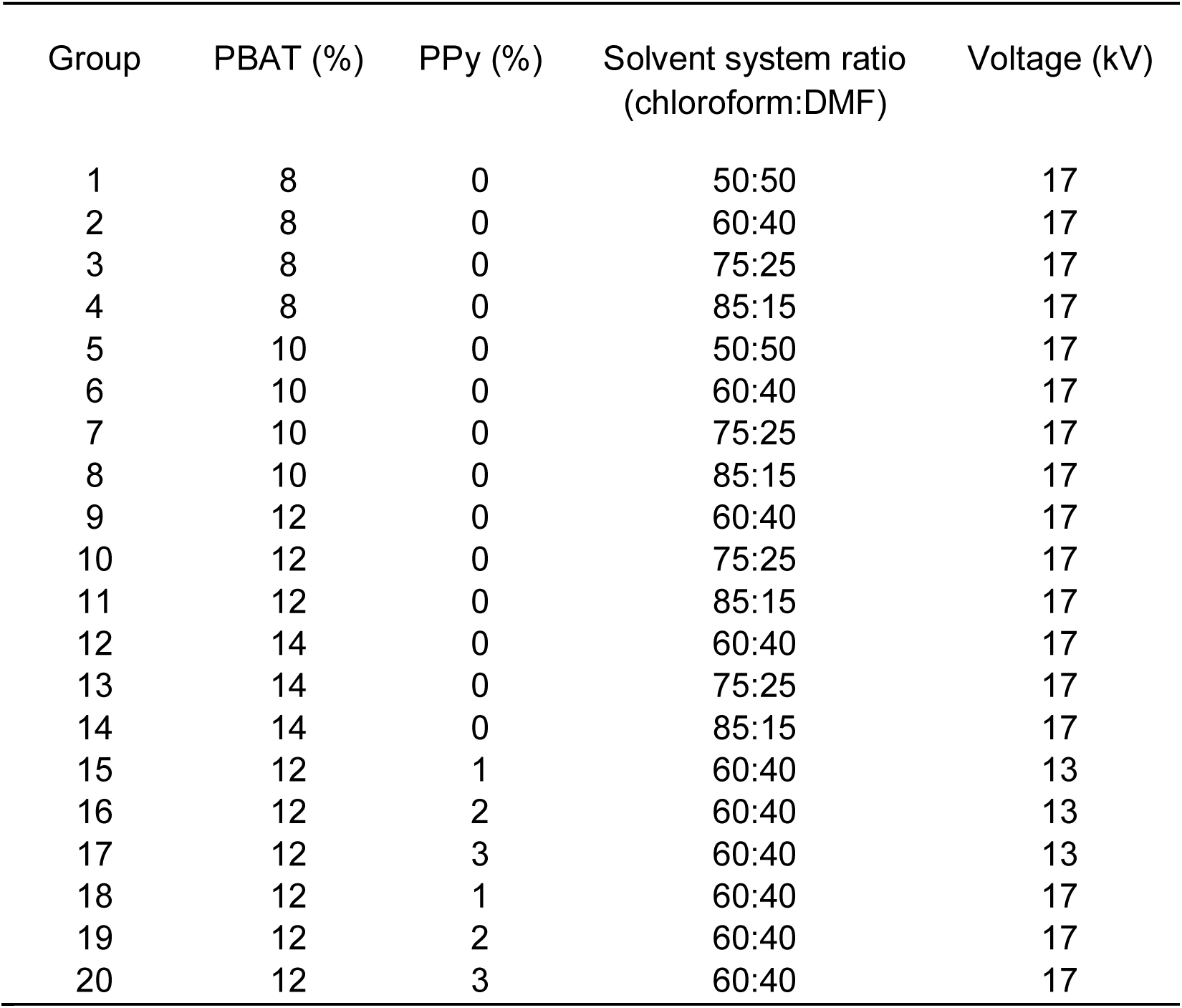
Percentage of PBAT, PPy and solvents, and voltage applied for electrospinning.

| Group | PBAT (%) | PPy (%) | Solvent system ratio<br>(chloroform:DMF) | Voltage (kV) |
| --- | --- | --- | --- | --- |
| 1 | 8 | 0 | 50:50 | 17 |
| 2 | 8 | 0 | 60:40 | 17 |
| 3 | 8 | 0 | 75:25 | 17 |
| 4 | 8 | 0 | 85:15 | 17 |
| 5 | 10 | 0 | 50:50 | 17 |
| 6 | 10 | 0 | 60:40 | 17 |
| 7 | 10 | 0 | 75:25 | 17 |
| 8 | 10 | 0 | 85:15 | 17 |
| 9 | 12 | 0 | 60:40 | 17 |
| 10 | 12 | 0 | 75:25 | 17 |
| 11 | 12 | 0 | 85:15 | 17 |
| 12 | 14 | 0 | 60:40 | 17 |
| 13 | 14 | 0 | 75:25 | 17 |
| 14 | 14 | 0 | 85:15 | 17 |
| 15 | 12 | 1 | 60:40 | 13 |
| 16 | 12 | 2 | 60:40 | 13 |
| 17 | 12 | 3 | 60:40 | 13 |
| 18 | 12 | 1 | 60:40 | 17 |
| 19 | 12 | 2 | 60:40 | 17 |
| 20 | 12 | 3 | 60:40 | 17 |

### Characterization of the surface tension of the solution

The characterization of the solution was performed by means of a tensiometer (Krüss EasyDyne K20) in an environment with controlled humidity and temperature, at 50% and 20 °C, respectively. For the measurements, solutions of PBAT (8%, 10%, 12% and 14 % m/v,) were prepared using chloroform and N, *N-*dimethylformamide as solvents, varying their proportions (Table 1).

### Electrospinning process

Electrospinning was carried out using a high voltage source (Spellman, CZE1000R) fixed distance of 10.0 cm between the needle tip (18 G, 1.2 mm) and the collector, and a solution flow rate of 1 mL h^-1^ (KDS 100). The temperature and humidity were carefully controlled at 23 °C and 30 %, respectively. The best parameters for the electrospinning process were investigated by varying the PBAT concentration, the applied voltage, the solvent system ratio and the PPy content (Table 1).

### Characterization by Scanning Electron Microscopy (SEM)

For this analysis, the samples were fixed in aluminum stubs by means of a carbon adhesive tape. Prior analysis, the samples were covered with a thin layer of gold, via sputtering technique (Emitech K550X). Micrographs with magnifications at 2000 and 5000 were obtained.

### Animals

The maintenance and care of the mice were performed in accordance with international standards on the use of laboratory animals. The protocols were reviewed and approved by the Committee of Ethics in the Use of Laboratory Animal from UNIFESP (CEUA 2101180516). C57/Bl6 mice from the Laboratory for Animal Experimentation (INFAR-UNIFESP) were kept in isolated units under SPF (specific pathogen free) sanitary conditions, at 20 ± 2 °C, relative humidity of 50 % and circadian cycle of 12 h interval (light/dark). Every effort was made to minimize animal suffering and reduce the number of animals used.

### Bone marrow mesenchymal stem cells isolation and culture

Bone-marrow-derived mesenchymal stem cells (BM-MSCs) were isolated from mice bones as previously described (17). Briefly, mice were euthanized by cervical dislocation, the tibias and femurs were dissected, and their epiphyses were cut. With a syringe needle 26 G (13 mm / 24.5 mm, BD Biosciences), 2 mL of DMEM (Dulbecco's Modified Eagle's Medium, Invitrogen) was injected at one end. The bone marrow was removed by flushing DMEM through the bone. The cell suspension was centrifuged at 400 × g for 10 min at room temperature. The precipitate was suspended in DMEM with 10 % fetal bovine serum (FBS; Cultilab), 1 % L-glutamine (Invitrogen) and 1 % penicillin / 1 % streptomycin (GibcoBRL). Cells were counted in a Neubauer chamber, diluted in the same medium at a density of 5 × 10^6^ cells / mL, and incubated at 37°C with 5 % CO_2_ atmosphere. The exchange of culture medium was performed every 72 h to remove nonadherent hematopoietic cells. When the cultures reached 90 % confluence, subcultures were performed using trypsin (Cultilab) in 10 mM PBS. After the second subculture, adherent cells were considered MSC as described in the literature (17, 18).

### Cell Viability Assay

BM-MSCs viability was assessed by MTT (3-(4, 5-dimethylthiazolyl-2)-2, 5-diphenyltetrazolium bromide) assay. Cells were cultured in 24-well plates for 24 h in the presence and absence of nanofibers immersed in the medium. At the end of this period 270 μL of culture medium and 30 μL of MTT solution (5 mg / L) were added to each well and incubated for 3 h at 37°C. After this period, MTT was aspirated and the reaction product, formazan salt, was dissolved by the addition of 180 μL of dimethylsulfoxide (DMSO, MP Biomedicals) in each well. The plate was shaken for 15 min, the content was transferred to a 96-well plate and the optical density was read at 540 nm on an ELISA plate reader (Labsystems Multiskan MS).

### Culture of mouse neuroblastoma cell line Neuro2a

Neuro2a cells were cultured according to the recommended protocol from ATCC (American Type Culture Collection). Briefly, cells were cultured in a 75 cm^2^ culture flask with 15 mL of culture medium composed of high glucose DMEM, 1% penicillin / streptomycin, 1 % glutamine and 10 % FBS. Cells were kept at 37 °C and 5 % CO_2_. At 60% confluence cells were trypsinized in the ratio of 1:5.

### Neuro2a differentiation protocol

Neuro2a cells are able to differentiate into neurons by decreasing FBS concentration from 10% to 0.5% and by adding 10 mM retinoic acid (RA) (19). To evaluate cell differentiation, Neuro2a cells were plated either on PBAT, PBAT/PPy 1%, 2%, 3%, or on coverslips, and incubated with regular culture medium (high glucose DMEM, 10 % FBS). After 24 h, culture medium was removed and differentiation medium (high glucose DMEM, 0,5% FBS, 10 mM RA) or regular culture medium, for undifferentiation control. After 72 h cells were fixed with 4 % paraformaldehyde and prepared for imunocytochemistry or Scanning Electronic Microscopy (SEM).

### Immunostaining

Neuro2a cells were grown either on coverslips, laminin-coated nanofibers (50 μg / mL; Sigma Aldrich), or on uncoated nanofibers for 96 h. After this period, cells were fixed with 4% paraformaldehyde, permeabilized with 0.1 % Triton X-100 (Sigma Aldrich), and immunollabeled with anti-Tubulin beta-3 (Tubbeta3) (Merck Millipore) followed by incubation with fluorophore-conjugated secondary antibody (Alexa 488 or 594) plus DAPI to stain DNA. Coverslips were assembled with Fluoromount G (Electron Microscopy Sciences) and analyzed using an inverted confocal microscope (Leica Microsystems). Image overlays were generated by ImageJ software.

### Measurement of size and number of Neuro2a neurites

Immunostained Neuro2a were photographed using inverted confocal microscopy. Neurite length was defined as the distance between the center of the cell soma and the tip of its longest neurite. Only neurites emerging from an isolated cell, but not from a clump of cells, were considered, and they should not contact other cells or neurites. Cell projections were considered neurites to be counted and measured when size ≥ 10 μm. For each experimental group, the lengths of neurites from 10 randomly selected cells from each of 10 photos for each group were measured using ImageJ software and the mean values were calculated according to a protocol in the literature (20).

### Analysis of cell adhesion by SEM

Neuro2a adhesion on nanofibers (PBAT, PBAT/PPy 1%, 2% and 3%) composites was evaluated after the differentiation period. Cells attached to the substrate were fixed with a 4 % paraformaldehyde and dehydrated in a graded ethanol solution series (30–100 %) for 10 min each. The drying stage used a 1:1 solution of ethanol with hexamethyldisilazane (HMDS), and the samples were dried with pure HMDS at room temperature. After deposition of a thin gold layer, the specimens were photographed using an XL30 FEG (Mira, Tescan) scanning electron microscope.

### Statistical Analysis

The statistical significance of the results was analyzed using Student’s t-test or one-way ANOVA. The results correspond to the average ± standard error, and results were considered statistically significant if p < 0.05.

## RESULTS

### Surface tension and SEM of the polimeric fibers

Surface tension was reduced with the increase of chloroform in the solutions, independently of the percentage of PBAT (Table 2). This factor can be associated with the solution density, as the increase in density resulted in lower surface tension values. Since the density of chloroform is 1.49 g / mL and the density of DMF is considerably lower (0.994 g / mL), the use of higher proportions of chloroform provided lower values of surface tension, closer to the density of chloroform.

**Table 2:** Density and surface tension of the soiutions at different proportions of soivents and concentration of PBAT.

| PBAT | Chloroform:DMF | Density (g mL<sup>-1</sup>) | Surface Tension (mN m<sup>-1</sup>) |
| --- | --- | --- | --- |
| 8% | 50:50 | 1.23 | 31.0 |
| 8% | 60:40 | 1.28 | 30.0 |
| 8% | 75:25 | 1.36 | 29.0 |
| 8% | 85:15 | 1.41 | 28.2 |
| 10% | 50:50 | 1.23 | 30.8 |
| 10% | 60:40 | 1.29 | 30.3 |
| 10% | 75:25 | 1.37 | 29.2 |
| 10% | 85:15 | 1.42 | 28.4 |
| 12% | 50:50 | 1.55 | - |
| 12% | 60:40 | 1.28 | 30.5 |
| 12% | 75:25 | 1.35 | 29.3 |
| 12% | 85:15 | 1.40 | 28.8 |
| 14% | 50:50 | - | - |
| 14% | 60:40 | 1.27 | 30.9 |
| 14% | 75:25 | 1.35 | 29.8 |
| 14% | 85:15 | 1.39 | 29.1 |

We analyzed the morphology of the electrospun PBAT fibers obtained from solutions prepared with the different chloroform/DMF ratios using SEM and identified the occurrence of defects known as “beads” (Figure 1). We observed that 8% PBAT solutions were unable to produce continuous fibers, producing agglomerated granules, with the formation of “bead-on-string” fibers coming off the beads, but very small in length and thin (Figure 1a), flattened regions, which may arise from the non-evaporation of the solvent to the target (Figure 1b,c). A greater agglomeration and thickening of the flattened regions were observed (Figure 1b), whereas these phenomena were more scattered, with the decrease in the amount of DMF in the solution (Figure 1c). Increased proportion of chloroform produced fibers with larger amount of granules, but it was possible to detect evidence of fiber formation (Figure 1d).

**Figure 1:**
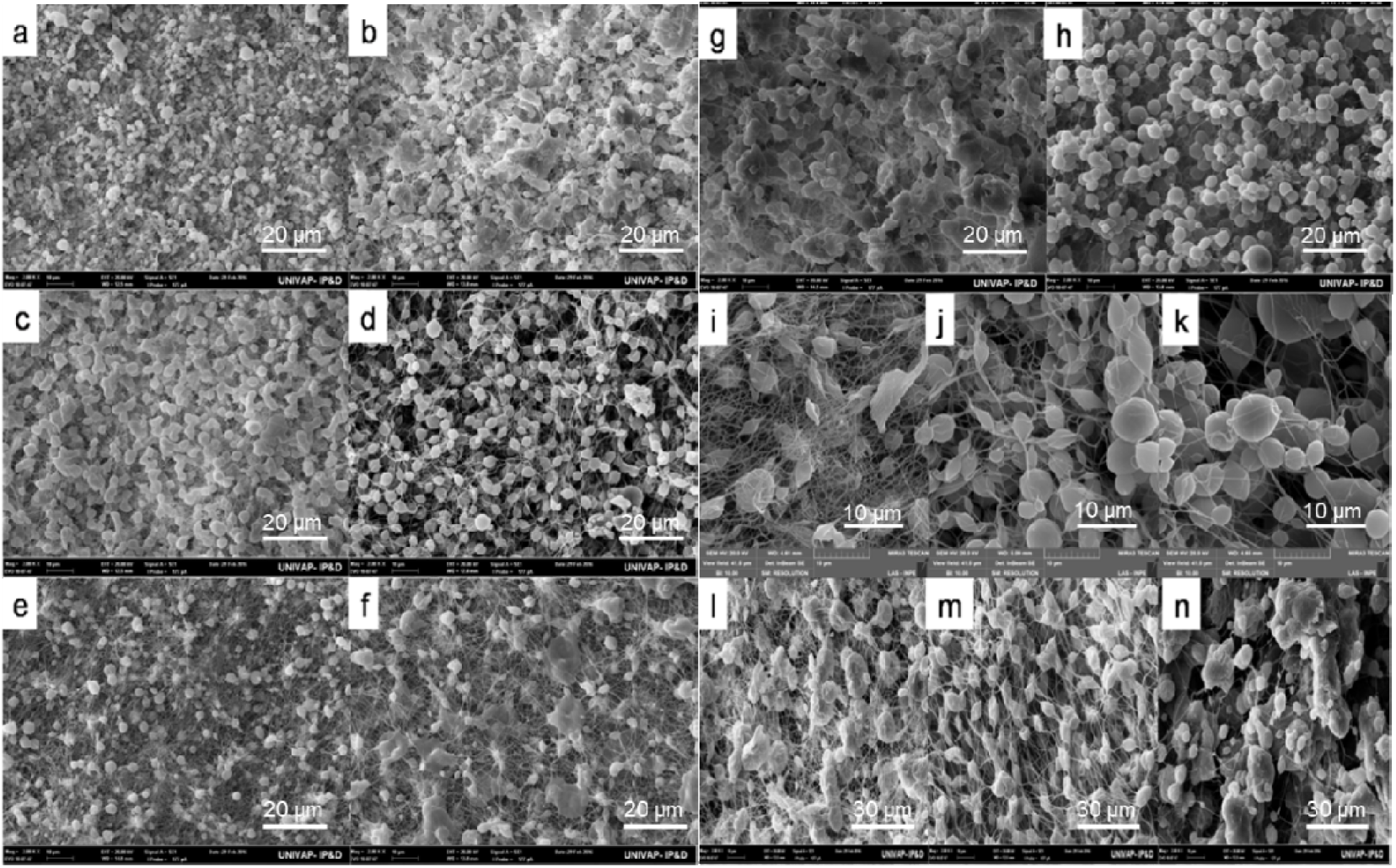
Scanning Electron Microscopy (SEM) of PBAT 8 % (a) 50:50, (b) 60:40, (c) 75:25, (d) 85:15; PBAT 10 % (e) 50:50, (f) 60:40, (g) 75:25, (h) 85:15; PBAT 12 % (i) 60:40, (j) 75:25, (k) 85:15; PBAT 14 % (l) 60:40, (m) 75:25, (n) 85:15.

A large number of beads can still be observed in fibers produced by 10 % PBAT, some of the polymer solutions were able to produce continuous fibers, which were sharper in the samples containing less chloroform, although we observe the formation of imperfect fibers with a great number of beads (Figure 1e). The presence of beads more widely scattered in 10 % PBAT with the formation of membranes from the incomplete evaporation of the solvent, was observed when chloroform:DMF proportion was 60:40 (Figure 1f). Increasing proportion of chloroform intensified the presence of thicker membranes with no significant fiber formation (Figure 1g). When the proportion of solvents was 85:15, the films disappeared, and numerous beads with few fibers filled with imperfections were formed (Figure 1h).

The analysis of 12 % PBAT samples, with different solvent ratios, indicated that the sample prepared using chloroform:DMF at 60:40 showed the best result among all samples prepared (Figure 1i). Although the presence of some beaded regions is still observed, it is possible to notice reduced imperfections in relation to the other groups and a denser fiber web. Fiber formation was reduced and the beads became more numerous, consequently not favorable for cell experiments, when the solvent proportion was either 75:25 or 85:15 (Figure 1j,k).

Samples prepared with 14 % PBAT, produced with different solvent proportion 60:40 (Figure 1l); 75:25 (Figure 1m), and 85:15 (Figure 1n) showed a viscosity limit. The incorporation of higher amounts of the polymer produced fibers with more imperfections, in the form of granules along the fibers, due to viscosity. The beads became more scattered and smaller with solvent proportion 75:25 (Figure 1m), whereas with 85:15 we observed very low incidence of fiber formation with the intensification of bead formation, giving rise to large polymeric clusters probably due to the inability to form a stable electric field during the electrospinning process (Figure 1n).

After evaluating the best parameters for obtaining electrospun PBAT mats, we started the production of PBAT/PPy fibers (PPy 1–3%; Figure 2). The applied voltage during the electrospinning process is a very important factor, since it is responsible for generating an electric field strong enough to break the surface tension of the solution (23). We explored voltages 13 kV and 17 kV to produce PBA/PPy fibers. Although presenting relatively higher standard deviation, samples electrospun at 17 kV showed better conformation, reducing the presence of beads at 1 % and 3 % of PPy (Figure 2d,f), as well as higher fiber spacings, which is a very important factor for cell adhesion, since the spacing between the fibers favors the anchoring of the cells and facilitates fluid flow (23). Therefore, for all cell experiments, we electrospun scaffolds at 17 kV.

**Figure 2:**
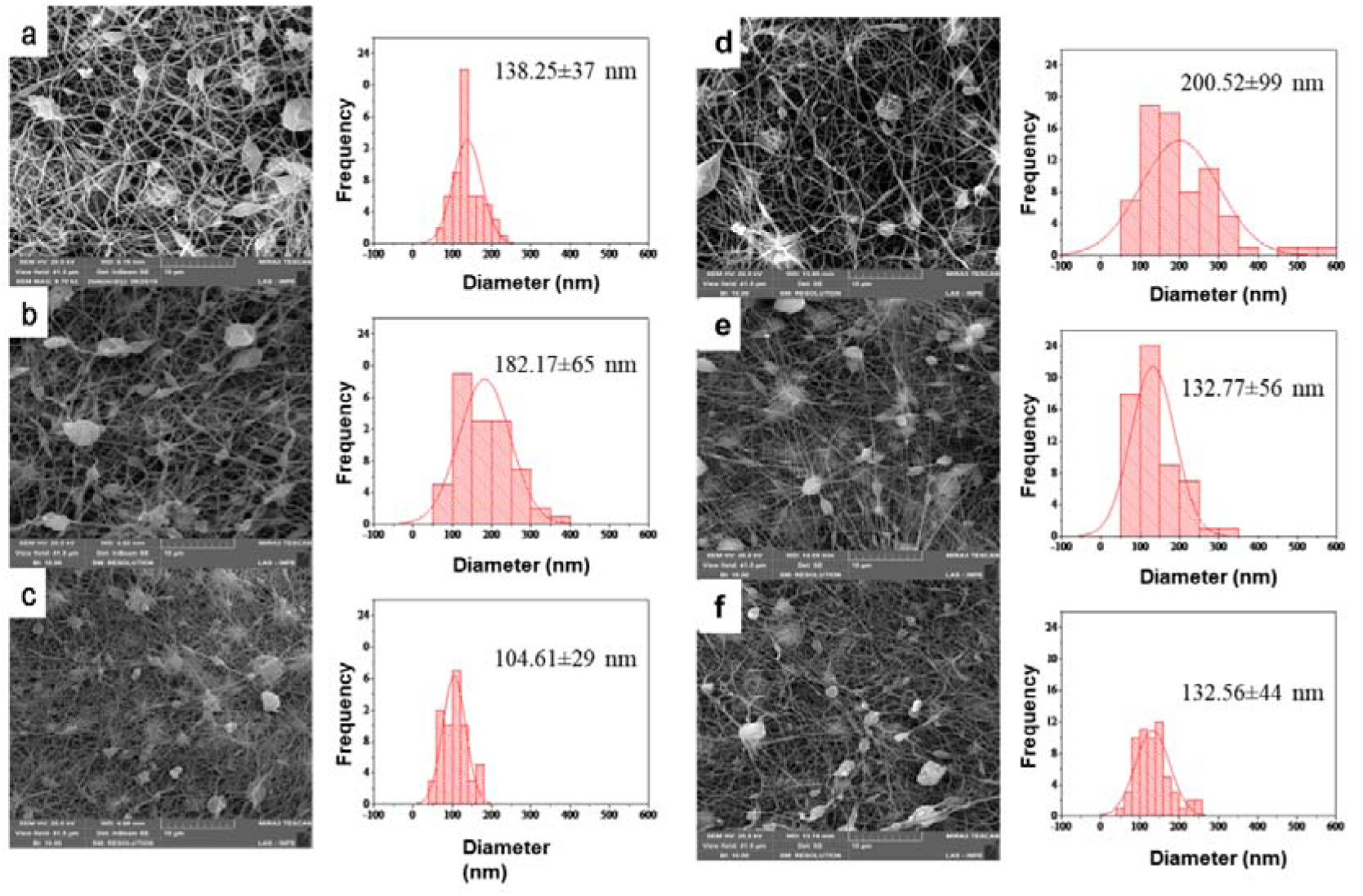
Scanning Electron Microscopy (SEM) refer to PBAT/PPy samples obtained using 13 kV and 17 kV, with Gaussian distribution: (a) 13 kV PBAT/PPy1%, (b) 13 kV PBAT/PPy 2%, (c) 13 kV PBAT/PPy3%, (d) 17 kV PBAT/PPy1%, (e) 17 kV PBAT/PPy2% and (f) 17 kV PBAT/PPy3%.

### PBAT/PPy polymeric fibers do not affect BM-MSCs viability and adhesion

We used mice BM-MSCs to test for biocompatibility of PBAT and PBAT/PPy prior to the use with the neuroblastoma cell line Neuro2a, because primary cell culture, i.e. cells that are isolated from tissues and cultured, tend to be more sensitive to death than cell lineages. We cultured MSCs on PBAT/PPy2%, measured the amount of live cells after 48 h, and did not observe differences when compared to cells cultured on glass coverslips (control) (Figure 3a), indicating that PBAT/PPy2% did not affect cells viability in 48 h. We also aimed to analyze if BM-MSCs adhered and extended protrusions when grown on PBAT/PPy2%, as they would do when cultured on plastic or glass. Using SEM, we observed that cells spread and extended cellular protrusions, exhibiting normal BM-MSC morphology (Figure 3b-e), suggesting PBAT/PPy2% did not impair cell adhesion.

**Figure 3:**
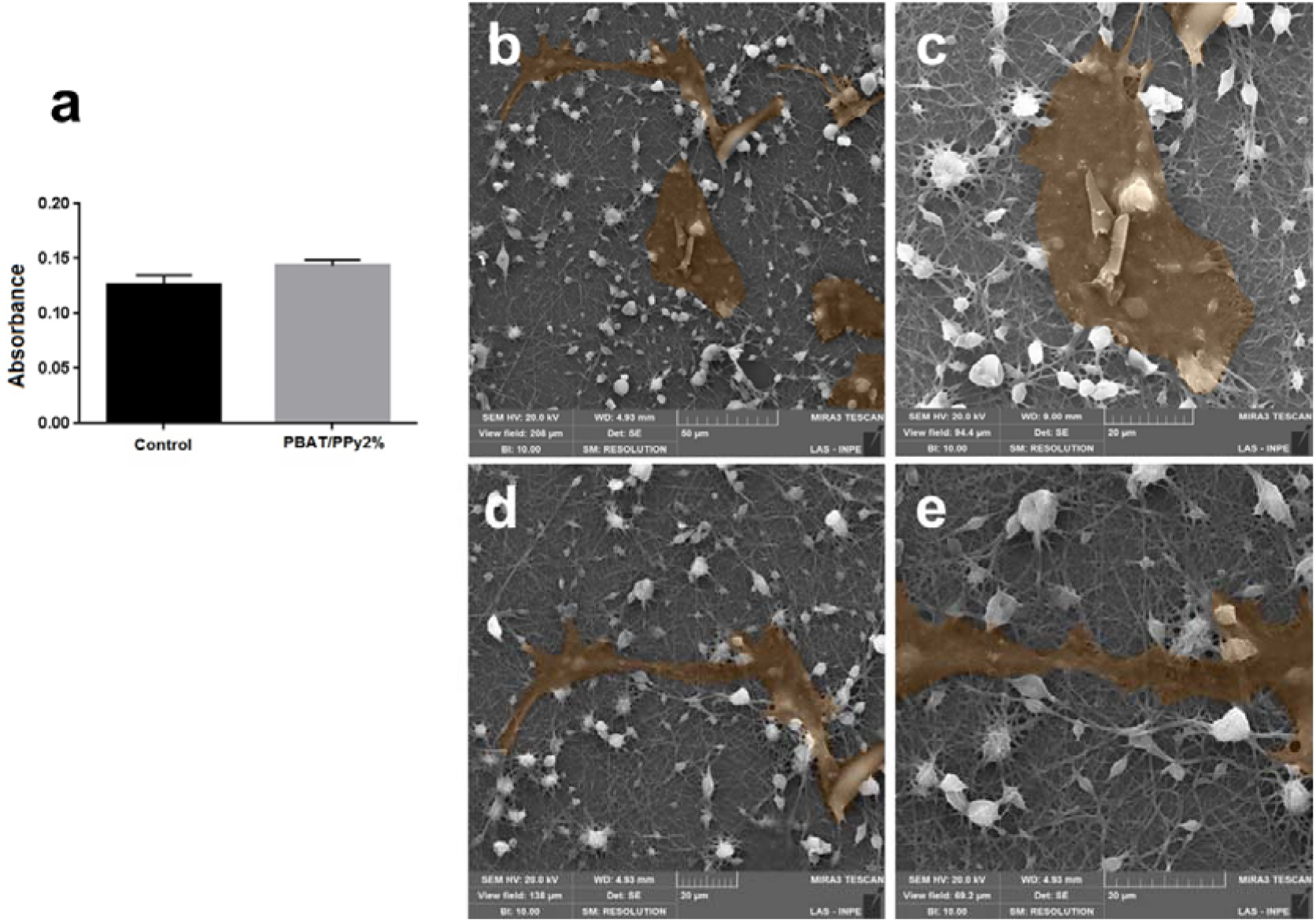
MTT assay absorbance values for Control and PBAT/PPy2% groups when measured by spectrophotometer (A). SEM images of BM-MSCs adhered on the fibers surface of PBAT/PPy2% for 48h (B-E). Student’s t test, considering significant if p < 0.05.

### PBAT/PPy polymeric fibers do not impair Neuro2a neuronal differentiation

Neuro2a cells were stimulated to differentiate for 3 days *in vitro* by treatment with RA and reduction of FBS concentration from 10 % to 0.5 %. Surprisingly, Neuro2a cells cultured on PBAT/PPy2%, and treated with RA in low serum (0.5 % FBS) developed more neurites, that were also longer, when compared to the same conditions for cells grown on fibers composed of PBAT only (Figure 4).

**Figure 4:**
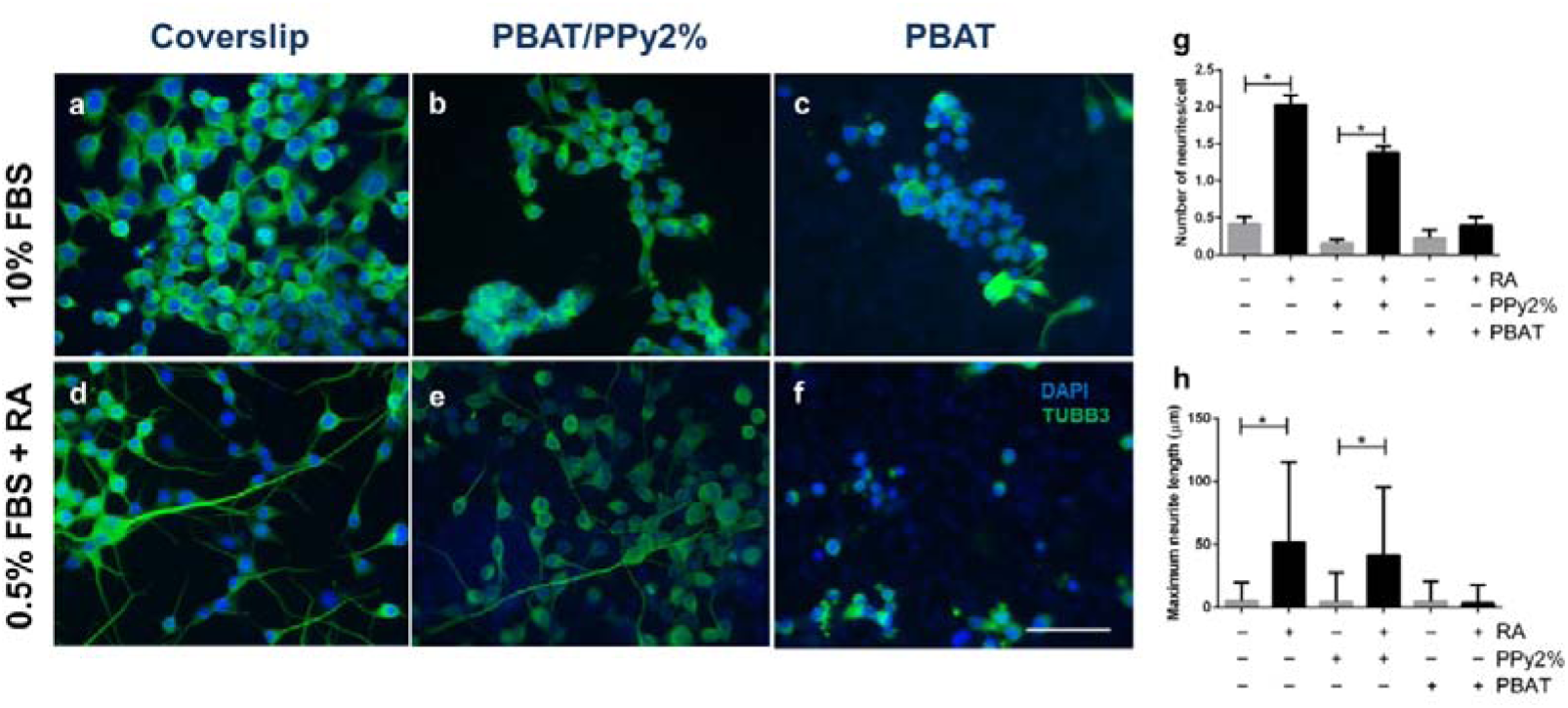
Neuro2a cells grown on coverslips (A and D), PBAT / PPy2% (B and E) and PBAT (C and F). Marker for mature neuron (Tubulin beta 3; TUBB3; green) and nuclear staining (DAPI; blue). Cells were cultured with DMEM + 10 % FBS in A, B and C. Cells were stimulated to differentiate with DMEM + 0.5 % FBS + retinoic acid (RA) in D, E, and F. Scale bar: 100 μm. (G) Number of neurites per cell. (H) Maximum neurite length per cell. One-way ANOVA, Tukey's multiple comparisons test (*p < 0.05; **p < 0.01).

In order to decrease hydrophobicity of PBAT, a characteristic that could impair Neuro2a differentiation, we cover PBAT fibers with a biomolecule that would increase PBAT hydrophilicity. For that purpose, we chose the extracellular matrix protein laminin. Laminins are a major component of the basal lamina, a protein network present in several tissues and organs, that serves as support for cell adhesion, migration and differentiation (24). The hydrophilic characteristics of laminin plus its role as an extracellular matrix component that binds to members of the integrin family of cell adhesion receptors should enhance cell adhesion as well as increase the number and length of neurites if hydrophobicity was responsible for inhibition of Neuro2a differentiation. Indeed, coating PBAT scaffolds with laminin increased the number of neurites per cell when the culture was treated with RA (Figure 5g) when compared to untreated cells. Interestingly, the number and length of neurites extended by Neuro2a cultured on laminin-coated PBAT scaffolds and treated with RA were similar to those observed for PBAT/PPy2% (Figure 5h). Decreasing PPy concentration to 1 % or increasing it to 3 % did not improve both parameters, suggesting that 2 % of PPy in PBAT was the best combination to ensure neurite extension.

**Figure 5:**
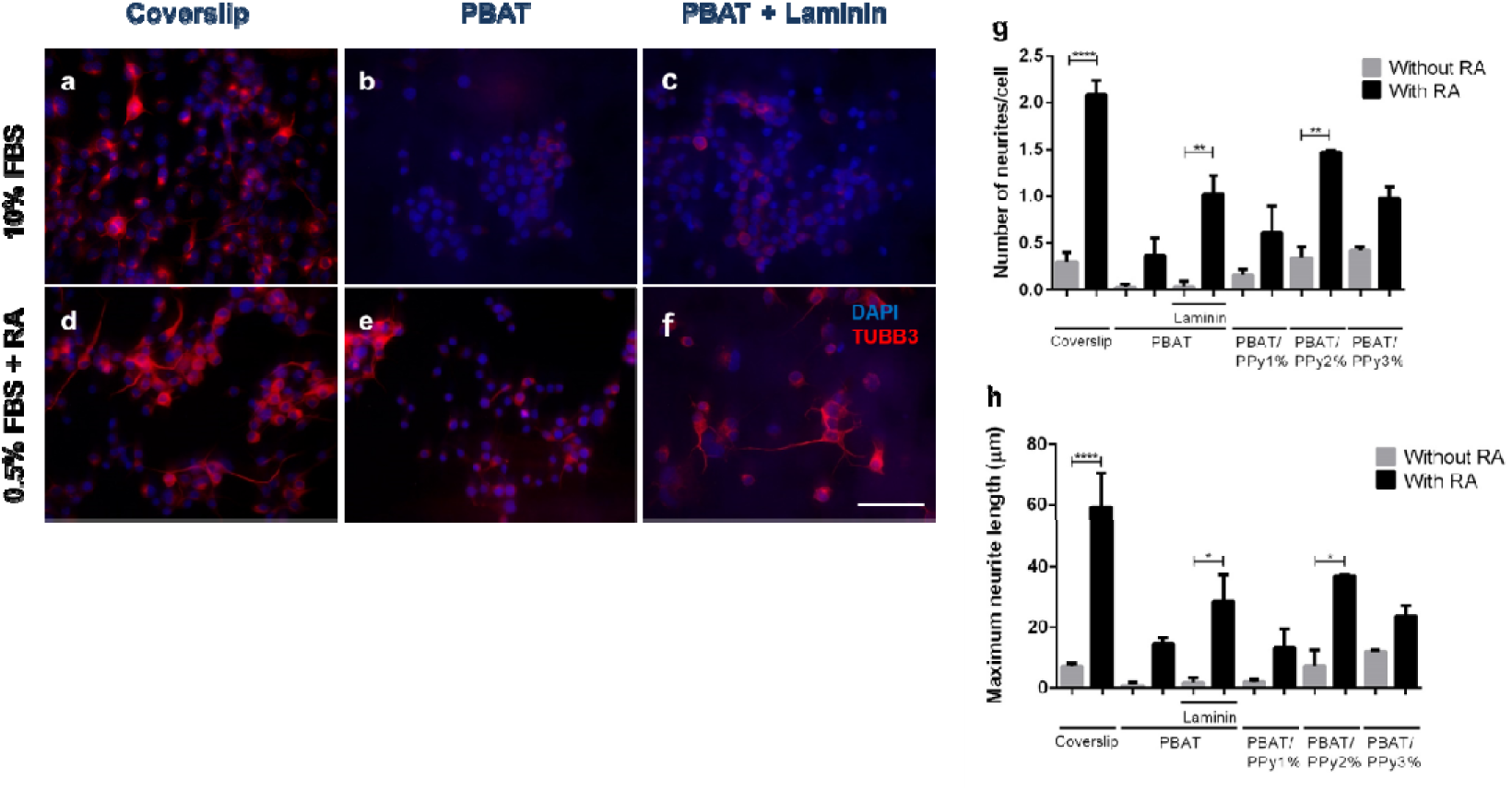
Neuro2a ceiis grown on coversiips (A and D), PBAT (B and E) and PBAT + Laminin (C and F). Marker for mature neuron (Tubuiin beta 3; TUBB3; red) and nuciear staining (DAPI; biue). Ceiis were cuitured with DMEM + 10 % FBS in A, B and C. Ceiis were stimuiated to differentiate with DMEM + 0.5 % FBS + retinoic acid (RA) in D, E, and F. Scaie bar: 100 μm. (G) Number of neurites per ceii. (H) Maximum neurite iength per ceii One-way ANOVA, Tukey's muitipie comparisons test (*p < 0.05; **p < 0.01).

## DISCUSSION

Alzheimer’s Disease, stroke and traumatic brain injuries are neurodegenerative disorders and injuries that affect the central nervous system. Due to characteristics of the neural tissue, full regeneration of the central nervous system is still not possible (25, 26) and is been sought by different means, including stem cell therapies (27, 28), and engineered materials (29–31).

In this study, we present a methodology for the production of electrospun hybrid materials composed of PBAT and PPy, biodegradable and conductive polymers, respectively. We first analyzed the structures formed by the combination of the polymers, depending on concentration of PBAT and PPy, as well as the voltage applied to generate the fibers. Using SEM we observed presence of beads (Figure 1), that may occur due to a variety of reasons, such as charge density, applied voltage, viscosity and surface tension of the solution (32). The formation of thin fibers, mainly at the nanoscale, can be attributed to the low viscosity of the polymer solutions, which facilitates deformation and consequently stretching during the electrospinning process. However, the instability of the jet elongation increases with the deformation of low viscosity polymer solutions, making it difficult to produce nano and ultrathin fibers free of beads (33). Samples containing smaller amounts of polymer showed a lot of beads, due to high proportion of chloroform, and the solutions that fit these characteristics also showed lower viscosity and surface tension (Table 2), which is in agreement with what have described (34–36). The sample of 12 % PBAT with proportion of solvent 50:50 was not favorable for electrospinning, due to the high viscosity. Highly viscous solutions are not favorable for electrospinning, as well as high amounts of the auxiliary solvent, which impair the total evaporation of the solvent during electrospinning (23).

Using SEM we observed that 12 % PBAT solubilized in chloroform:DMF (60:40) presented better conformation in relation to the other proportions studied. The most suitable values of density and surface tension were 1.280 g / mL and 30.5 mN / m, which were intermediate values when compared to the others (Table 2). Based on the arguments of De Brito et al, we argue that very low surface tensions promoted instability in the electrospinning process (23).

Addition of PPy to the polymer solution reduced the amount of beads formed (Figure 2). This factor is related to the conductive nature of PPy, which facilitates the breakdown of the surface tension, stretching the solution with consequent formation of the Taylor Cone (22). Although with relatively higher standard deviations, samples electrospun at 17 kV showed better conformation, reducing the presence of beads (1 % and 3 % PPy), as well as higher fiber spacings, which is a very important factor for cell adhesion tests, since spacing between fibers favors anchoring of cells and facilitates fluid flow (23).

Reduction of imperfections can be associated to the increase in diameter of the fibers produced at 17 kV. This factor can be related to phenomena already reported in the literature (23). However, the reduction of the granules is also associated with the modification of the voltage, since higher voltages have been shown to be more effective. This fact is explained by Costa et al. citing that the stability of the process is directly associated to the applied voltage (22). However, even with beads, the fibers presented a cylindrical shape, demonstrating that the electrospinning parameters, such as moisture and applied tension, were adequate, otherwise the fibers would have flattened conformation, resembling tapes.

Our ultimate goal with this study was to produce fibers that would allow for neural cells to attach and spread. We have previously studied PLA fibers as scaffolds to support attachment and growth of BM-MSCs that were transplanted to mouse brain submitted a stroke model (37). MSCs produce and secrete a variety of biological factors, such as growth and survival factors, cytokines, neurotrophins, among others (38). Transplantation of MSCs improves recovery in animal models of cerebral ischemia by mechanisms that may include neuroprotection, induction of axonal sprouting, and neovascularization (39). In this study we used BM-MSCs to test the cytotoxicity of the fibers.

PBAT and PBAT/PPy are described as non-cytotoxic materials (13, 40, 41). We cultured BM-MSCs on PBAT/PPy2%, and observed that the fibers were biocompatible (Figure 3). It is known that the cytotoxicity of a biomaterial is affected by several factors such as its surface topology and hydrophilicity (42). As we observed that PBAT/PPy2% did not affect cell viability we assume that it is suitable for biomedical use in accordance with ISO–10993-5.

We investigated the morphology of BM-MSC cultured on PBAT/PPy2%, and observed that the cells exhibited plasma membrane protrusions with maintenance of normal MSC morphology (Figure 3), suggesting they are well adhered to the fibers. It is known that tridimensional scaffolds favor cell adhesion due to the fact that their arrangement is similar to the extracellular matrix (43). Typical characteristics of cytotoxic cellular changes were not observed. Similar results were obtained from our group regarding cell adhesion on PBAT scaffolds, showing the initial phase of cell adhesion with cells spreading with no preferential direction, resulting in the formation of a monolayer after 3 days in culture (13, 44).

To study the compatibility of PBAT/PPy fibers with neural cells we chose the neuroblastoma cell line Neuro2a, cells used in various neural applications including the study of neurite outgrowth (45). When Neuro2a are cultured in 10 % FBS they stay in an undifferentiated state. If concentration of FBS is decreased to 0.5 % and RA is added to the culture medium, cells differentiate in mature neurons that express tubulin beta3 and extend neurites (19). Our results showed that Neuro2a cultured on PBAT/PPy2% and treated with RA in 0.5 % FBS have more and longer neurites when compared to cells grown on PBAT only (Figure 4). The use of PPy combined with other degradable polymers such as PLCL, PLLA, PCL has already shown promising results regarding neuronal viability and neurite extension, especially when neuronal cells are electrically stimulated (30, 46, 47).

Another characteristic of a material reported to increase cell viability is surface wettability, the more hydrophilic a material is, the less cell death occurs (48). Additionally, increase in surface wettability increases the number and length of neurites (49). Therefore hydrophilicity of PPy used in PBAT/PPy2% could explain the increase in number and length of neurites observed as compared to neat PBAT, which is a hydrophobic polymer (50). In a previous work of our group we described the ability of PPy to induce osteoblastic differentiation, and a possible explanation relied on wettability properties (13). Therefore, the dynamic contact angle between a deionized water drop and the surface of PBAT and PBAT/PPy nanofibers was measured. We observed an advanced contact angle (ACA, at 2 min) of 84° for PBAT/PPy, while for PBAT the ACA was higher (115°). This result showed that PPy improved the wettability of PBAT scaffolds. Few authors observed a relationship between surface hydrophobicity and cell spreading (51, 52).

To verify if the hydrophobicity of PBAT impairs Neuro2a differentiation, we coated PBAT fibers with a biomolecule that could change its hydrophobicity, the extracellular matrix protein laminin. Laminins are a major component of the basal lamina, a protein network present in several tissues and organs, that serves as support for cell adhesion, migration and differentiation (24). The hydrophilic characteristics of laminin plus its role as an extracellular matrix component that binds to cell adhesion receptors (integrins) should improve cell adhesion with consequent differentiation, if hydrophobicity was responsible for impairment of Neuro2a differentiation. Indeed, coating PBAT scaffolds with laminin increased de number of neurites per cell when the culture was treated with RA (Figure 5). Interestingly, the number and length of neurites extended by Neuro2a cultured on laminin-coated PBAT scaffolds and treated with RA were similar to those observed for PBAT-PPy2% (Figure 5). Decreasing to 1% or increasing to 3% did not improve both parameters, suggesting that 2% of PPy in PBAT was the best combination to ensure neurite extension.

Finally, in this work we intended to develop a composite for future *in vivo* neuroregeneration applications without using any element isolated from animals that could possibly generate an unwanted immune response (53). The results, as a whole, showed that the developed composite PBAT/PPy improved interactions with cells when compared to PBAT alone. This study opened possibilities to generate new conductive biomaterials for neural applications based on a still poorly exploited thermoplastic copolymer (PBAT/PPy). Other studies are still necessary to determine the electrical properties of PBAT/PPy fibers, morphological improvement of the material, evaluation of gene expression when cells are cultured on the fibers, as well as *in vivo* experiments, in order to explore the full potential use of PBAT/PPy fibers in regenerative medicine, specially in neurology.

